# Weighted RSA: an improved framework on the perception of audio-visual affective speech in left insula and superior temporal gyrus

**DOI:** 10.1101/2020.08.31.276485

**Authors:** Junhai Xu, Haibin Dong, Fei Guo, Zeyu Wang, Jianguo Wei, Jianwu Dang

## Abstract

Being able to accurately perceive the emotion expressed by the facial or verbal expression from others is critical to successful social interaction. However, only few studies examined the multimodal interactions on speech emotion, and there is no consistence in studies on the speech emotion perception. It remains unclear, how the speech emotion of different valence is perceived on the multimodal stimuli by our human brain. In this paper, we conducted a functional magnetic resonance imaging (fMRI) study with an event-related design, using dynamic facial expressions and emotional speech stimuli to express different emotions, in order to explore the perception mechanism of speech emotion in audio-visual modality. The representational similarity analysis (RSA), whole-brain searchlight analysis, and conjunction analysis of emotion were used to interpret the representation of speech emotion in different aspects. Significantly, a weighted RSA approach was creatively proposed to evaluate the contribution of each candidate model to the best fitted model. The results of weighted RSA indicated that the fitted models were superior to all candidate models and the weights could be used to explain the representation of ROIs. The bilateral amygdala has been shown to be associated with the processing of both positive and negative emotions except neutral emotion. It is indicated that the left posterior insula and the left anterior superior temporal gyrus (STG) play important roles in the perception of multimodal speech emotion.

## Introduction

In daily communication, the processing of emotion perception is multimodal. We can know the emotion state of speakers not only through facial expressions, but also from body movements, voice rhythms and other information. Successful social interactions require an accurate understanding of others’ emotions, intentions and thoughts. In particular, emotion expression provides critical information about the emotion state or surroundings of the expresser. These signals can help guide behavior and even have survival values in some situations. Therefore, being able to accurately perceive the emotions of others from their facial expressions and voice messages is very important in social interactions. Previous studies on emotion perception usually use unimodal stimuli, mainly focusing on emotion perception and recognition of facial or auditory stimuli. Although few studies focused on the representation of audio-visual emotions, instrumental sounds and emotional music were mostly collected as the auditory stimuli. Compared to music or instrumental sounds, affective speech is closer to normal social interactions. However, researches on speech emotion are limited, and their findings were inconsistent. Therefore, it remains unclear how speech emotion is perceived in the brain with multimodal stimuli.

One specific network on the unimodal emotion processing is observed by lots of researches. In terms of visual modality, studies have shown that the fusiform gyrus, right superior temporal sulcus, inferior frontal gyrus and amygdala were significantly more activated by emotional facial expressions than neutral ones (Kesler-West et al. 2001). One latest study on emotion perception test using static facial expressions found that the emotion assessment test was associated with the functional connectivity strength from the fusiform gyrus to the frontal lobe and insula, indicating the role of these brain regions in the emotion perception test (Bae et al. 2019). Other studies using static facial emotion stimuli have found that the amygdala plays a significant role in emotion processing (Fitzgerald et al. 2006; Kugel et al. 2008). One of our previous studies suggested that the left central posterior gyrus could abstractly represent emotion expressed by face, body and whole person regardless of modality information (Cao et al. 2018). With dynamic facial expression stimuli, a neural network including the right amygdala, left globus pallidum and medial prefrontal cortex were observed to be more activated when an angry or happy face than a neutral one was viewed (MORIOKA et al. 2010). For the auditory modality, increased activations were found in the amygdala, hippocampus, and temporal pole when listening to unpleasant music compared to pleasant music (Koelsch et al. 2006). Moreover, one study have used different musical instruments to express different types of emotions and found that the insula could identify specific emotion categories ignoring the type of musical instruments being played (Sachs et al. 2018). In order to get closer to the daily communication environment, more studies adopted emotion rhythms as stimuli, and found that bilateral superior temporal gyrus and sulcus might be involved in the processing of emotion rhythm information (Leitman et al. 2010; Dara 2012; Ethofer et al. 2012). These studies often focused on unimodal and didn’t involve in the multimodal interactions of emotion even though they pointed out the role of some brain regions in emotion processing. It remain unclear how the emotion information is perceived by human brain in response to multimodal stimuli.

Many studies have used combined visual and auditory emotion information to further explore the mechanism of cross-modality representation of emotions. For the audio-visual integration of emotion, a large number of studies showed that the superior temporal gyrus and superior temporal sulcus played an important role in the integration and control of audio-visual emotion information (Kreifelts et al. 2007; Robins et al. 2009; Park et al. 2010; Müller et al. 2012; Hagan et al. 2013). However, little research has been conducted on emotion perception in audio-visual modality. The brain regions that had a greater activation have been explored by emotional stimuli in audio-visual modality versus neutral, suggesting that bilateral superior temporal gyrus showed a significant effect in emotion perception (Robins et al. 2009). Moreover, multi-sensory emotion perception was believed to be represented by enhanced activation of brain regions revealed by multi-sensory integration, mainly including the superior temporal gyrus and superior temporal sulcus (Jessen and Kotz 2015). A recent study using meta-analysis of 18 neuroimaging studies identified a core audio-visual emotion processing network, including the right posterior superior temporal gyrus, the left anterior superior temporal gyrus, the right amygdala and the thalamus (Gao et al. 2019). The important role of the right superior temporal gyrus in facial perception of different emotion valence was further reinforced by examining the neural mechanism of emotion perception in visual and auditory modalities via the multivoxel pattern analysis (Zhang et al. 2019). These studies have expanded our understanding of emotional speech perception. However, the exact neural network involved in speech emotion perception was not consistent, and which brain regions might be involved in multimodal affective speech perception remained to be verified.

Traditional univariate approaches have been used in a series of neuroimaging studies to explore the representation mechanisms of multimodal affective information, such as the generalized linear model (GLM). By modeling and analyzing the experimental conditions, the GLM can obtain the activation information of each voxel, and use statistical analysis to report the significantly activated voxel, which is likely to cause the loss of fine-grained pattern information (Haynes and Rees 2006; Norman et al. 2006). Currently, more advanced methods, such as multivoxel pattern analysis (MVPA) and representational similarity analysis (RSA), make it possible to analyze the activation pattern information throughout the whole-brain level. RSA is a computational method for multi-channel measures of neural activity proposed by Kriegeskorte et al., university of Cambridge, which can bridge the complexity of activation patterns between different brain-activity measurements and between subjects and species (Kriegeskorte et al. 2008). Compared with the univariate encoding models that predict each response channel separately, RSA focuses on the study of representational geometry, making the findings more intuitive (Xue et al. 2010; Devereux et al. 2013; Nili et al. 2014; Bracci et al. 2015). This study used hypothesis-driven RSA to analyze the correlation between neural representations in specific brain regions and models by constructing abstract emotion models. Most previous studies used RSA to compute the relationship between the representation of brain regions and a single model. In order to take all models into consideration, we creatively proposed a weighted RSA method, using weighted linear combinations of all abstract emotion models to construct the best fitted models of specific brain regions, so as to evaluate the contribution of each candidate model in the optimal model.

Previous studies have found that the amygdala is involved in emotion perception in visual or auditory modality (Fitzgerald et al. 2006; Koelsch et al. 2006; MORIOKA et al. 2010). Numerous studies have revealed the role of the superior temporal gyrus in multimodal affective perception and integration (Robins et al. 2009; Leitman et al. 2010; Park et al. 2010; Jeong et al. 2011; Müller et al. 2012; Hagan et al. 2013; Jessen and Kotz 2015; Gao et al. 2019; Zhang et al. 2019). The insula has been pointed out to be involved in the perception and experience of emotions (Duerden et al. 2013), and the insula cortex have been confirmed to be necessary and sufficient platform for human emotions, which was actually the only neural source of emotion experience (Damasio et al. 2013). In addition, some studies have revealed that the inferior parietal lobule might be related to the cross-modal decoding of emotion, indicating that it might also be involved in emotion perception (Kim et al. 2017). In this study, we took the amygdala (AMG), insula, inferior parietal lobule (IPL) and superior temporal gyrus (STG) as regions of interest (ROI) to further investigate their role in cross-modal interactions. Based on findings in the literature on unimodal emotion perception and recent studies on dynamic audio-visual emotional cues, it is hypothesized that the insula and STG are involved in perception of speech emotion in audio-visual modality.

Previous studies have revealed the processing mechanism of emotion in facial expressions, music or instruments. However, it is still unclear how the speech emotion is perceived in visual and auditory modalities. In this study, we made an event-related design to explore the brain regions capable of perceiving emotional and neutral stimuli in all modalities, when participants experience five stimuli (anger, sad, neutral and joy) expressed in three modalities (visual, auditory and audio-visual). After modeling the data using GLM, RSA based on ROIs was used to examine the correlation between abstract emotion models and neural representation patterns in specific brain regions. Then a novel weighted RSA was proposed to evaluate the contribution of each candidate model to the best fitted model. Moreover, the searchlight-based RSA was conducted to further explore the brain region that was significantly associated with a specific model. Finally, the conjunction analysis of emotion was used to further determine the candidate areas involved in emotion perception. We hope to prove our experiment with variety of advanced approaches in different aspects in order to provide evidence for the perception mechanism of multimodal speech emotion in human brain.

## Materials and Methods

### Participants

Twenty-five healthy volunteers were recruited in this study (ten females mean age 23.3 ± 1.40 years, range from 21 to 26 years). All subjects were right-handed, had normal or corrected-to-normal vision, and had no history of neurological or psychiatric problems. For the data quality control, nine subjects were excluded for further analysis. This study was carried out in accordance with the recommendations of Institutional Review Board (IRB) of Tianjin Key Laboratory of Cognitive Computing and Application, Tianjin University. All subjects gave written informed consent in accordance with the Declaration of Helsinki. After the experiment, all subjects will be paid accordingly.

### Experiment Stimuli

The stimuli in this study came from Geneva Multimodal Emotion Portrayals (GEMEP) (Bänziger et al. 2011), which is a video dataset, performed and recorded by 10 professional actors (5 males and 5 females). The stimuli adopted in the experiment included anger, sad, neutral and joy emotions, and each emotion had five short sentences. These sentences were expressed by two actors (one male, one female) with facial expressions, in a total of 40 video clips (4 emotions × 5 sentences × 2 actors). The Adobe Premiere Pro CC 2014 was used to separate the picture track and audio track of the video, and extract the dynamic facial expression and sound of each video respectively, with duration of 2 seconds, for the single visual or auditory stimuli. The audio-visual condition reintegrated the clipped dynamic pictures and sounds to present visual and auditory emotional information at the same time.

### Procedure

The fMRI experiment contained three runs, which were emotion recognition tasks of facial expressions, emotional rhythm as well as consistent audio-visual emotion. Among them, the first run was emotion recognition of dynamic facial expression, which was designed to recognize emotions through facial expressions regardless of audio information. The second run was designed to identify the emotions by the speech rhythm ignoring the picture information. The third run was the consistent study of audio-visual emotion, in which the same emotion was used as the visual and auditory modalities. The three runs were similar, with different stimulus materials. Fig. 1 showed the design exemplar of the first run.

**Fig 1.**
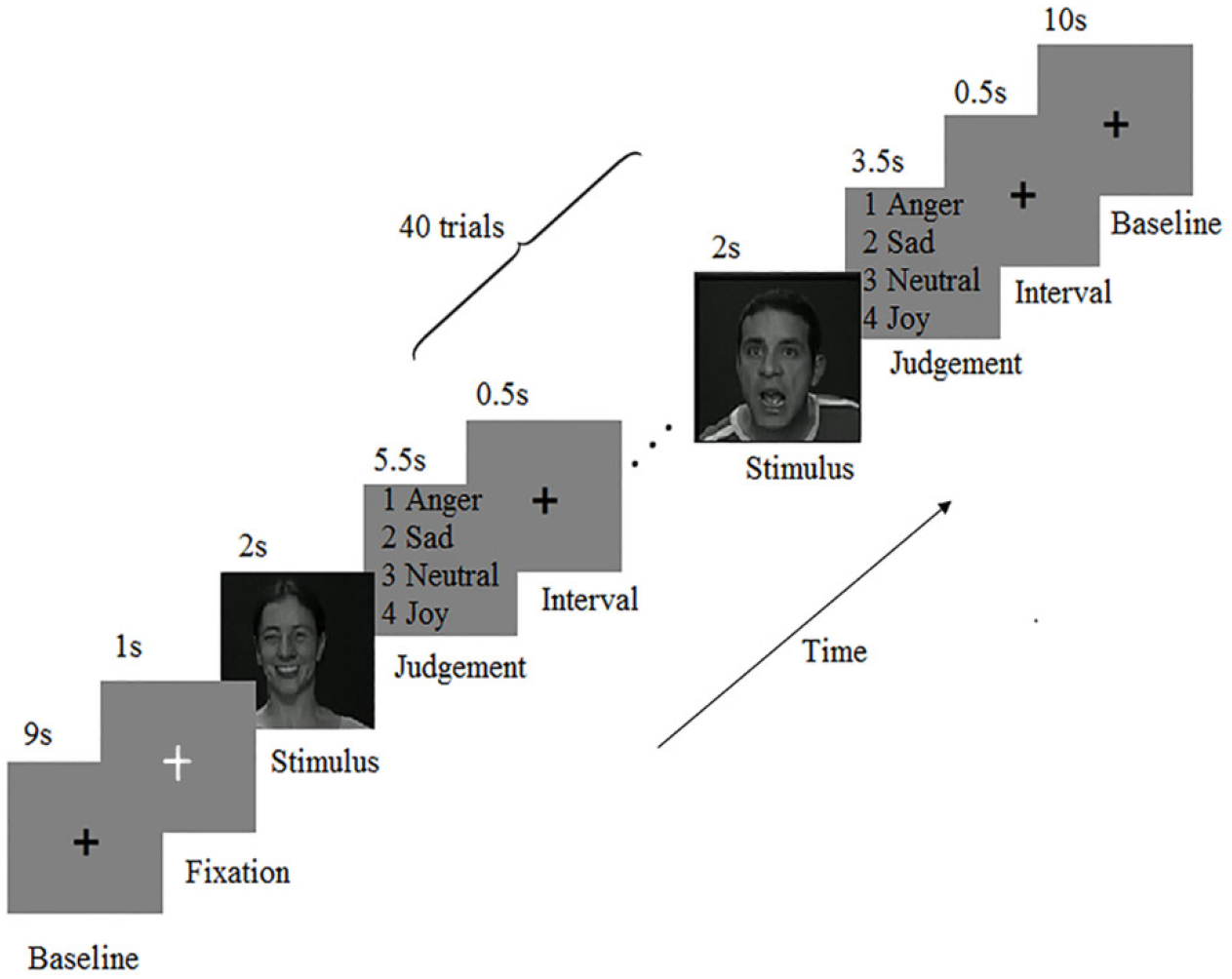
The design of the first run. The beginning displayed a black cross for 9s and a white cross for 1s. Then a sequence of stimuli was presented, each consisting of a stimulus for 2s and an emotion recognition (anger, sad, neutral and joy) as well as an interval for 4 to 6 seconds. The number of stimuli was 40 (4 emotions × 10 trials for each emotion), and there was a fixed interval for 10s in the end.

An event-related design was used in this experiment. There was a fixed interval of 10s at the beginning and end of each run. Each run had 40 trials, including 10 trails for each condition of anger, sad, neutral and joy. In each trial, the stimuli displayed for 2 seconds, and the inter stimulus interval was 4 to 6 seconds with an average interval of 5 seconds. In each run, the order of the stimuli was pseudo-randomly.

### Data Acquisition

The imaging data was collected at Tianjin Huanhu Hospital using a 3.0T Siemens magnetic resonance scanner and an eight-channel head coil. Functional images were acquired by an echo-planar imaging (EPI) sequence, with the parameters as follows: TR (repetition time) = 2000 ms, TE (echo time) = 30 ms, voxel size = 3.1 × 3.1 × 4.0 mm^3^, slices = 33, slices gap = 0.6 mm, slices thickness = 4.0 mm, FA = 90 degree, FOV = 200 × 200 mm^2^. In addition, a high-resolution anatomical image (T1-weighted image) was acquired using a three-dimensional magnetization-prepared rapid-acquisition gradient echo (3D MPRAGE) sequence, with the following parameters: TR = 1900 ms, TE = 2.52 ms, TI = 1100 ms, FA = 9 degree, Voxel size = 1 × 1 × 1 mm^3^, FOV = 256 × 256 mm^2^. During the experiment, foam pads were used to reduce head movements and ear plugs were used to reduce scanner noises. A high-resolution stereo 3D glass of VisualStim Digital MRI Compatible fMRI system was required for visual stimuli required to wear and the electrostatic headphones were used for the auditory stimuli.

### Data Analysis

#### Data preprocessing

Data preprocessing was performed using the SPM12 toolkit (http://www.fil.ion.ucl.ac.uk/spm/software/spm12/) in Matlab software (The Math Works). The first five volumes of the functional data corresponding to the baseline of each run were discarded in order to eliminate the effect of unstable signal of the scanner at the beginning of experiment. The T1-weighted image was segmented into white matter, gray matter and cerebrospinal fluid (CSF). Functional images were mapped to the Montreal Neurological Institute (MNI) space after the operation of slice timing, realign and co-register. The standard of head movement is that the horizontal movement is less than 2mm and the rotation angle is less than 1.5 degree. All of the subjects met the standard in the experiment. Then the T1-weighted image was co-registered to the mean functional images for further normalization. Subsequently, the functional data were smoothed with a 6-mm full-width at half-maximum (FWHM) Gaussian filter aiming to improve the signal to noise ratio.

#### Generalized Linear Model (GLM)

After data preprocessing, the generalized linear model (GLM) was conducted to model the data of each subject. GLM is based on the assumption that the experimental data on each voxel (represented by Y) is a linear combination (represented by β) of unknown parameters (represented by x). The unknown parameters include the interested part (12 stimulus conditions, 3 modalities × 4 emotions), the uninterested part (6 head motion parameters) and residual (represented by *ε*). The general form of GLM is as follows:

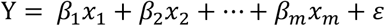

After modeling, the statistical analysis of experimental data was transformed into statistical inference of parameter beta, and the restricted maximum likelihood method was used to fit out the value of the parameter beta to minimize the sum of error.

#### Representational Similarity Analysis (RSA)

RSA can use the abstract models to interpret the representation of specific brain regions. Using the beta values obtained from GLM, RSA can calculate the correlation between the activation patterns of different stimulus conditions in specific brain regions, and then acquire the dissimilarity between different conditions, so as to obtain the representation dissimilarity matrices (RDMs) of brain regions. Meanwhile, the RDMs of specific model can be constructed using hypothetically driven RSA. Computing the correlation between the neural representation RDM of a specific brain region and the model RDMs can estimate whether the brain region contains the information expressed by the model. In this study, a computational framework was proposed with the RSA Toolbox from the aspect of ROIs and an exploratory “searchlight” analysis was designed to analyze the representation pattern of speech emotion in the human brain.

#### RSA based on ROIs

The steps of the RSA based on ROIs were as follows:

First, construct the neural representation RDMs. For a specific brain region, the beta values of all voxels for each stimulus condition obtained by GLM were extracted and expanded to a matrix of 12 × n (where 12 is the number of stimulus conditions and n is the number of voxels in the brain region). Then, for all voxels in this ROI, the Pearson correlation coefficients between each two conditions were calculated, and the dissimilarity values (1 minus Pearson correlation coefficients) were acquired. Thus, the data RDM of the brain region was constructed. Next, construct the candidate model RDMs. In this study, we constructed five emotion models (anger, sad, neutral, joy, and negative) and three modality models (visual, audio, and audio-visual) for the candidate model analysis. In the emotion models, the dissimilarity values between different modalities of the same emotion were set to 0 (for example, excited facial expressions and excited audio rhythm), and the dissimilarity degrees between other conditions were set to 1. The dissimilarity values between different emotions of the same modality were set to 0 (for example, excited facial expressions and sad facial expressions) and other parts of the RDMs were set to 1 in the modality models. The negative emotion model regarded anger and sadness as the same kind of emotion. Since the dissimilarity degrees on the diagonal were independent of the hypothesis, they were set to NaN and were excluded from the subsequent analysis.

Finally, make the statistical inference. For each ROI of each subject, the Kendal rank correlation coefficients between the data RDM and the RDMs of each model were calculated. In the group analysis, subjects were treated as random effects. The Kendall rank correlation coefficients were submitted to the unilateral Wilcoxon sign rank test to evaluate the contributions of the candidate models in explaining the neural representation patterns of a specific brain region.

#### Weighted RSA

To examine the contribution of the candidate models, we creatively proposed an approach of weighted RSA. The weighted linear sum of the 8 candidate models was used to fit one optimal model, and the mean square error between the optimal model and neural representation RDM of the brain region was minimized. By analyzing the weight of each candidate model in the fitted model, the contribution of the model could be estimated.

The detailed construction process of the weighted optimal model is as follows (Fig. 2). For the neural representation RDM of a specific brain region, it was expanded into a column vector with 144 elements, each of which represents a stimulus pair. For each element in the vector, the label was determined according to the criteria: if the value was less than 0.4, it was labelled as 0; the value greater than 0.6 was labelled as 2; if it was between 0.4 and 0.6, label it as 1. The Label vector corresponding to the element vector could be obtained. The function of the Label vector was to sort the elements with similar dissimilarity degrees to the same category and apply it to the generation of the optimal model. Then, the RDMs of the candidate models and a confound model were expanded to a matrix of 144 × 9. One of the row vectors of the matrix called “temp data” was extracted, and the label of the element in the data vector corresponding to that row could be obtained (for example, the label was 2). The row vectors in the matrix whose label were the same as the “temp data” were composed to a new matrix. Then a weight vector was initialized. In order to minimize the mean square error between the data vector and the column vector that was obtained by the product of the new matrix and the weight vector, continuous iterations were needed. A fitted element could be obtained by multiplying the temp data and the weight vector. By repeating the process for all stimulus pairs, the fitted vector could be acquired and the fitted model RDM could be obtained by transforming the vector to a square matrix.

**Fig 2.**
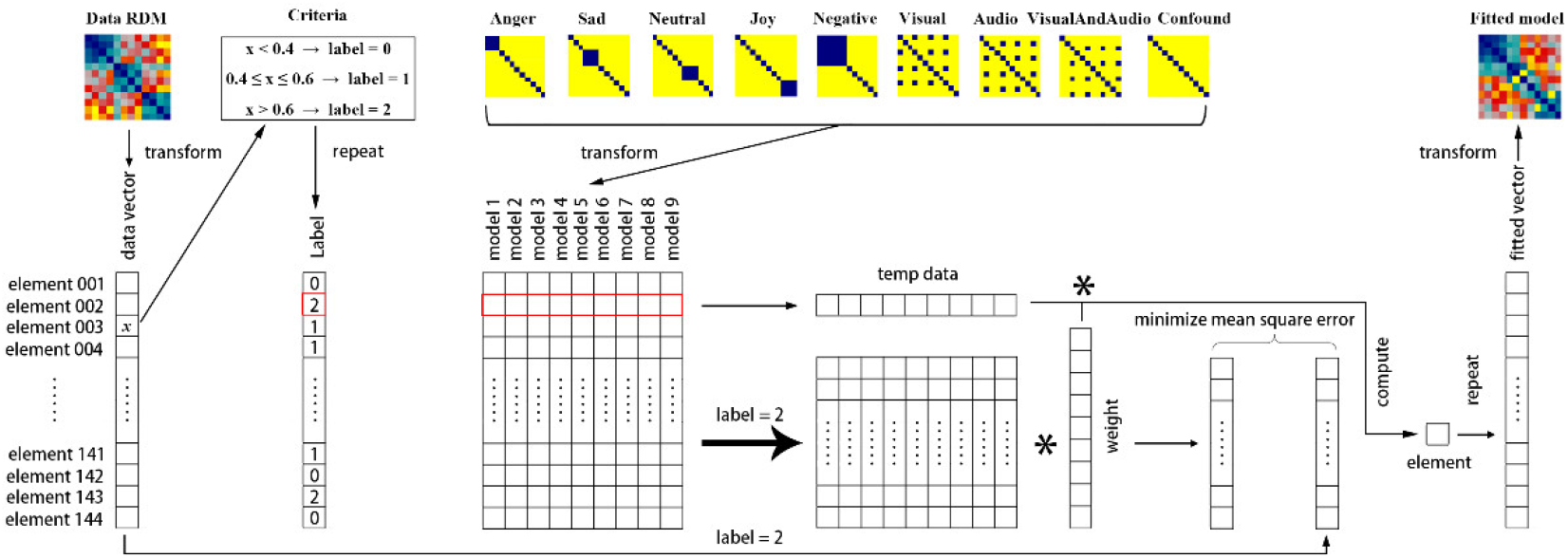
The construction process of weighted optimal model. There were three kinds of main procedures. First, the data RDM of a brain region was transformed to a vector and the Label vector could be obtained according to the criteria. Second, the candidate models were expanded to a matrix and the rows that had the same label with the temp data were combined to a new matrix. The weight vector could be obtained by minimizing the mean square error between the data vector and the product of the new matrix and the weight vector. Finally, multiplying the temp data and weight vector and repeating the process, the fitted model could be obtained by transforming the fitted vector to a square matrix.

#### Searchlight analysis

Brain regions that were significantly correlated with candidate models could be obtained in the whole brain by using searchlight analysis. In the searchlight analysis, for each voxel of each subject, the data in the neighborhood with a radius of 6 mm was extracted and was expanded in line. A matrix of 12 × n (where 12 is the number of stimulus conditions and n is the number of voxels in the searchlight) could be obtained for each searchlight step. Then, for each condition pair, Pearson correlation coefficients between the two conditions were calculated, and the 12 × 12 RDM of the searchlight was obtained. The Kendall rank correlation coefficient between the searchlight RDM and each candidate model RDM was calculated, which was assigned to the central element of the searchlight. The correlation coefficient could evaluate to what extent the neural representation pattern of the searchlight could be interpreted by the model. For each candidate model, the above process was repeated on all voxels of each subject, and the subjects’ whole brain correlation map (r-Map) was thus obtained. The group analysis was set with the subjects as random effects. For all candidate models, one-sample t-test was used to test the hypothesis of each voxel, and the significance value corresponding to the candidate model was obtained for each voxel, and then the whole brain significance map (p-Map) was obtained at the group level. Brain regions that were significantly correlated with the models were acquired with the threshold *p* = 0.01, corrected at the cluster level with the cluster size above 30 voxels.

#### Conjunction analysis of emotion

To further locate the regions that might be involved in emotion perception, three new contrasts were defined in the first level analysis for each subject, which were Anger > Neutral, Sad > Neutral and Joy > Neutral. Then, the results of all subjects were assessed in the second level group analysis, the significance level of which was set to *p* = 0.05 (FDR corrected) to obtain the activation maps of the three emotions versus neutral emotion. Finally, a conjunction analysis of (Anger>Neutral)∩(Sad>Neutral)∩(Joy>Neutral) was conducted based on the above results.

## Results

### RSA based on ROIs

Based on previous findings, the AMG, insula, IPL, and STG were initially defined to be the ROIs to further explore their roles in the perception of speech emotion in visual and auditory modalities. To precisely investigate the hemispheric effect on the representation of emotion, all ROIs were divided into two parts.

### Weight analysis of best fitted model

For each ROI, a best fitted model using the linear combination of the 8 candidate models was constructed to minimize the mean square error with the data RDM (Fig. 3). The weight analysis of emotion models revealed that compared with other brain regions, the negative model provided a small but significant contribution in the fitted results of the bilateral amygdala (left amygdala: Mean = 0.013, SD = 0.011; Right amygdala: Mean = 0.047, SD = 0.027), which indicated their roles in the processing of negative emotion. The joy emotion model had a significant contribution to the best fitted model of bilateral AMG, suggesting that they might also be associated with the processing of positive emotion.

**Fig 3.**
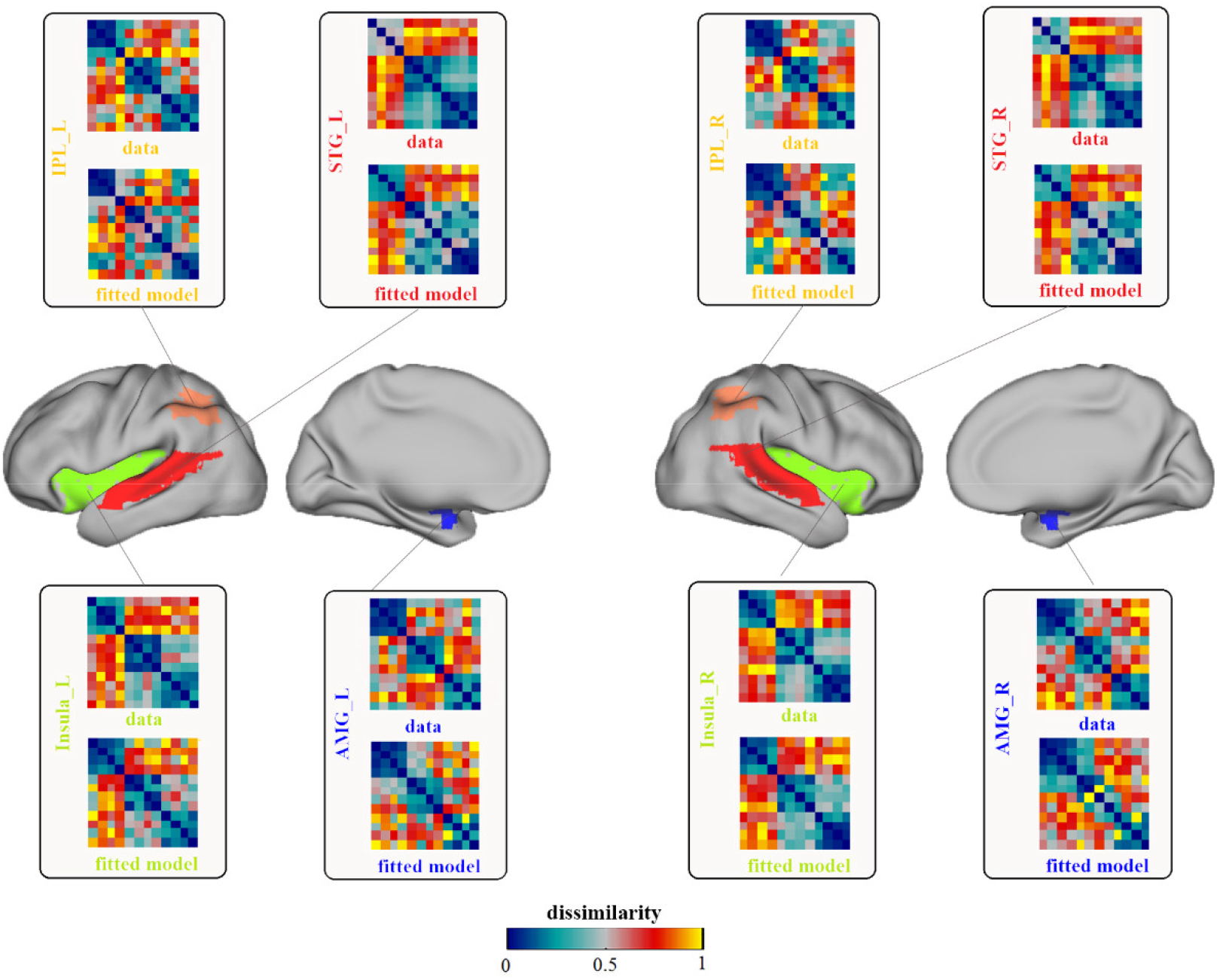
The results of data RDMs and fitted model RDMs on ROIs. Each pane showed the neural representation (data) RDM and the fitted model RDM of a specific brain region. The four panes on the left showed the results of left hemisphere and others showed the right hemisphere’s results. IPL, inferior parietal lobule; STG, superior temporal gyrus; AMG, amygdala; L, left; R, right.

The anger and joy emotion models shared significant contributions in all ROIs while the weights of sad and neutral models were relatively less. For the best fitted model of all brain regions, the weights of the neutral emotion model of the left AMG and the right insula were zero, while each single emotion model had a certain weight on other ROIs, indicating that they might be involved in the perception of speech emotion in different valences. For more details, please refer to Tab. 1.

**Tab. 1.** The weights of each model in fitted model (*p* = 0.05, FDR corrected)

| Region | Anger |  | Sad |  | Neutral |  | Joy |  | Negative |  |
| --- | --- | --- | --- | --- | --- | --- | --- | --- | --- | --- |
|  | Mean | SD | Mean | SD | Mean | SD | Mean | SD | Mean | SD |
| AMG_L | 0.404 | 0.230 | 0.056 | 0.118 | 0.000 | 0.000 | 0.208 | 0.103 | 0.013 | 0.011 |
| AMG_R | 0.376 | 0.202 | 0.044 | 0.108 | 0.108 | 0.144 | 0.175 | 0.105 | 0.047 | 0.027 |
| Insula_L | 0.307 | 0.243 | 0.008 | 0.012 | 0.043 | 0.018 | 0.036 | 0.134 | 0.001 | 0.001 |
| Insula_R | 0.266 | 0.306 | 0.042 | 0.127 | 0.000 | 0.000 | 0.127 | 0.246 | 0.000 | 0.000 |
| IPL_L | 0.217 | 0.130 | 0.051 | 0.041 | 0.005 | 0.004 | 0.062 | 0.082 | 0.000 | 0.000 |
| IPL_R | 0.283 | 0.170 | 1.13e-04 | 0.001 | 0.015 | 0.012 | 0.013 | 0.037 | 0.024 | 0.022 |
| STG_L | 0.259 | 0.346 | 0.020 | 0.018 | 0.002 | 0.002 | 0.085 | 0.192 | 0.000 | 0.000 |
| STG_R | 0.221 | 0.290 | 0.040 | 0.070 | 1.98e-04 | 0.001 | 0.069 | 0.141 | 0.000 | 0.000 |

SD, standard deviation; IPL, inferior parietal lobule; STG, superior temporal gyrus; AMG, amygdala; L, left; R, right.

### Statistical analysis

Neural representation RDM in each ROI was used to calculate the Kendall rank correlation coefficient with each model RDM, which were then examined with a unilateral sign rank (p set to 0.05, FDR corrected). We found that the data RDMs of all ROIs had no significant correlation with the sad emotion model. A significant correlation to the negative model was observed in the bilateral AMG, the values of which were greater than other emotion models. The correlation of joy model and bilateral AMG was also significant. These findings further indicated that the bilateral AMG might play an important role in the perception of positive and negative speech emotion in visual and auditory modalities. Bilateral AMG and neutral emotion model were not significantly correlated, suggesting that the bilateral AMG might not be associated with the processing of neutral emotion.

Bilateral insula, bilateral STG and left IPL showed significant correlations with all single emotion models except the sad model, which indicated their roles in the perception of emotions in different valences. No significant correlation with the neutral model was observed in the right IPL.

The statistical analysis results of all ROIs were shown in Fig. 4. It showed that the best fitted model was superior to any other models for all ROIs.

**Fig 4.**
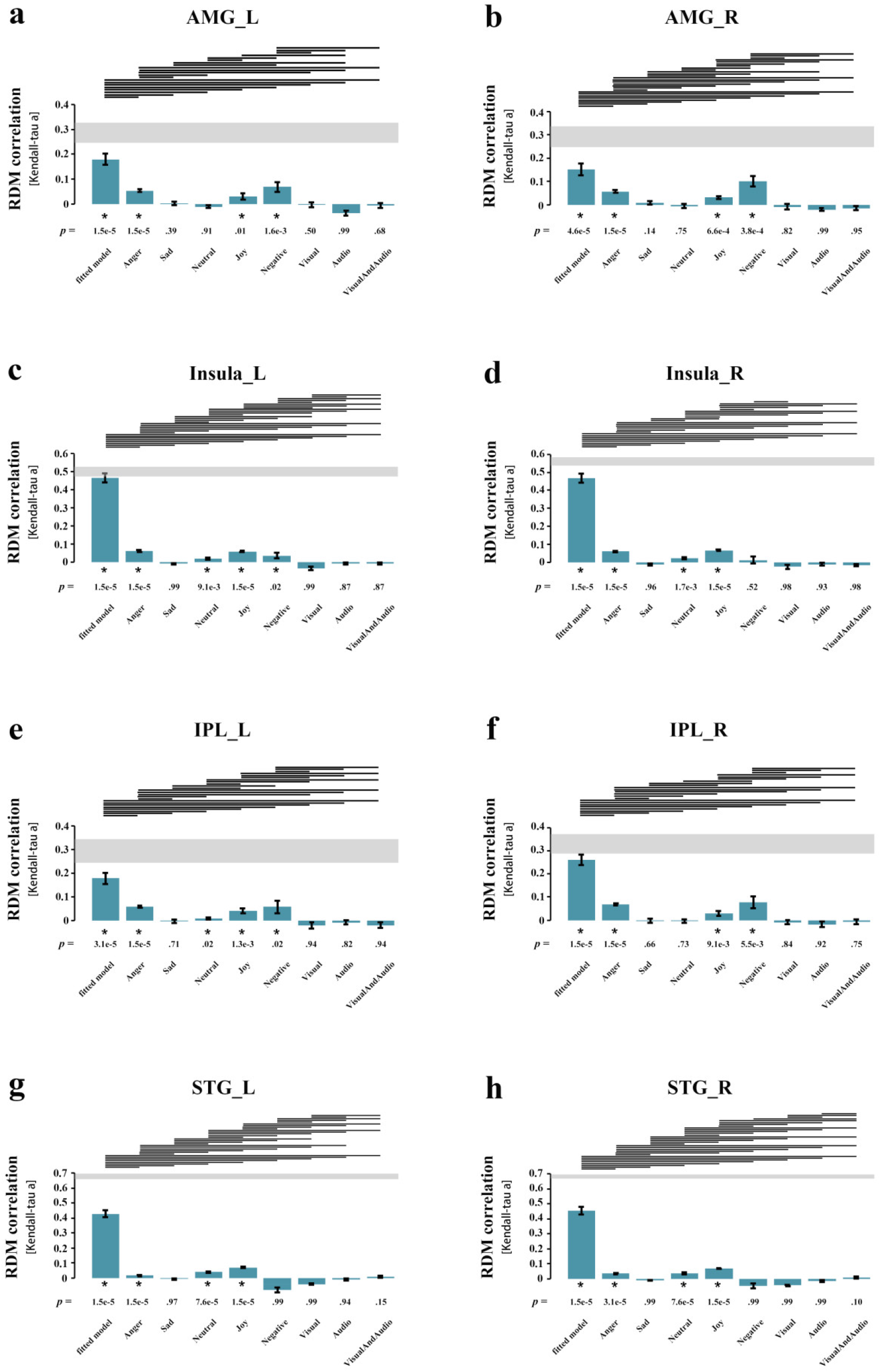
Statistical analysis results of ROIs. Each pane corresponded to the result of a ROI and showed the Kendall rank correlation coefficients between neural representation RDM of the ROI and the models. The models included the 8 candidate models and the best fitted model obtained by weighted RSA. P value stood for significance level and the asterisk represent the correlation was significant. The line segment meant the left model was superior to the right one. IPL, inferior parietal lobule; STG, superior temporal gyrus; AMG, amygdala; L, left; R, right.

### Searchlight analysis on the whole-brain level

The weighted RSA analysis showed that the weights of neutral emotion model were zero in the best fitted models of left AMG and right insula and the statistical analysis suggested that the correlation between the data RDM of right IPL and the neutral emotion model RDM was not significant. In order to explore whether these brain regions could perceive neutral emotion and promote the following analysis, a whole-brain searchlight analysis with a radius of 6mm was conducted for neutral emotion model. With a significant level of *p* = 0.01 (t-test) and cluster sizes above 30 voxels, brain regions that were significantly correlated with neutral emotion model were obtained.

As shown in Tab. 2, the whole-brain searchlight indicated that the correlation of bilateral IPL, bilateral insula and bilateral STG with neutral model reached the significant level (*p* = 0.01, t-test). Combined with the results of the statistical analysis, these brain regions were significantly correlated with all single emotion models except the sad emotion model, suggesting that they might be involved in the perception of speech emotion.

**Tab. 2.** The searchlight results of neutral emotion model (*p* = 0.01, t-test)

| Anatomical region | Hemisphere | Cluster size | MNI coordinates |  |  |
| --- | --- | --- | --- | --- | --- |
|  |  |  | x | y | z |
| STG | L | 598 |  |  |  |
| IPL | L | 171 | -51 | -6 | -24 |
| Insula | L | 60 |  |  |  |
| STG | R | 522 |  |  |  |
| Insula | R | 146 | 57 | 3 | -21 |
| IPL | R | 44 |  |  |  |

The cluster size indicated number of voxels; IPL, inferior parietal lobule; STG, superior temporal gyrus; L, left; R, right.

### Conjunction analysis on the emotion

To further explore if the IPL, insula and STG were really involved in the basic perception of speech emotion, a conjunction analysis of (Anger>Neutral)∩(Sad>Neutral)∩(Joy>Neutral) were conducted for all subjects’ results.

As shown in Fig. 5, anger, sad and joy emotion were significantly activated in the left posterior insula and left anterior STG versus neutral emotion (*p* = 0.05, FDR corrected, cluster size > 10). The results of weighted RSA showed that each single emotion model had a contribution to the best fitted model of left insula and left STG. Statistical analysis indicated that the neural representation RDMs of left insula and left STG were significantly correlated to all single emotion models except sad emotion model and whole brain searchlight analysis further revealed their role in processing neutral emotion. Considering the above results, we concluded that the left posterior insula and left anterior STG were involved in the perception of speech emotion in visual and auditory modalities.

**Fig 5.**
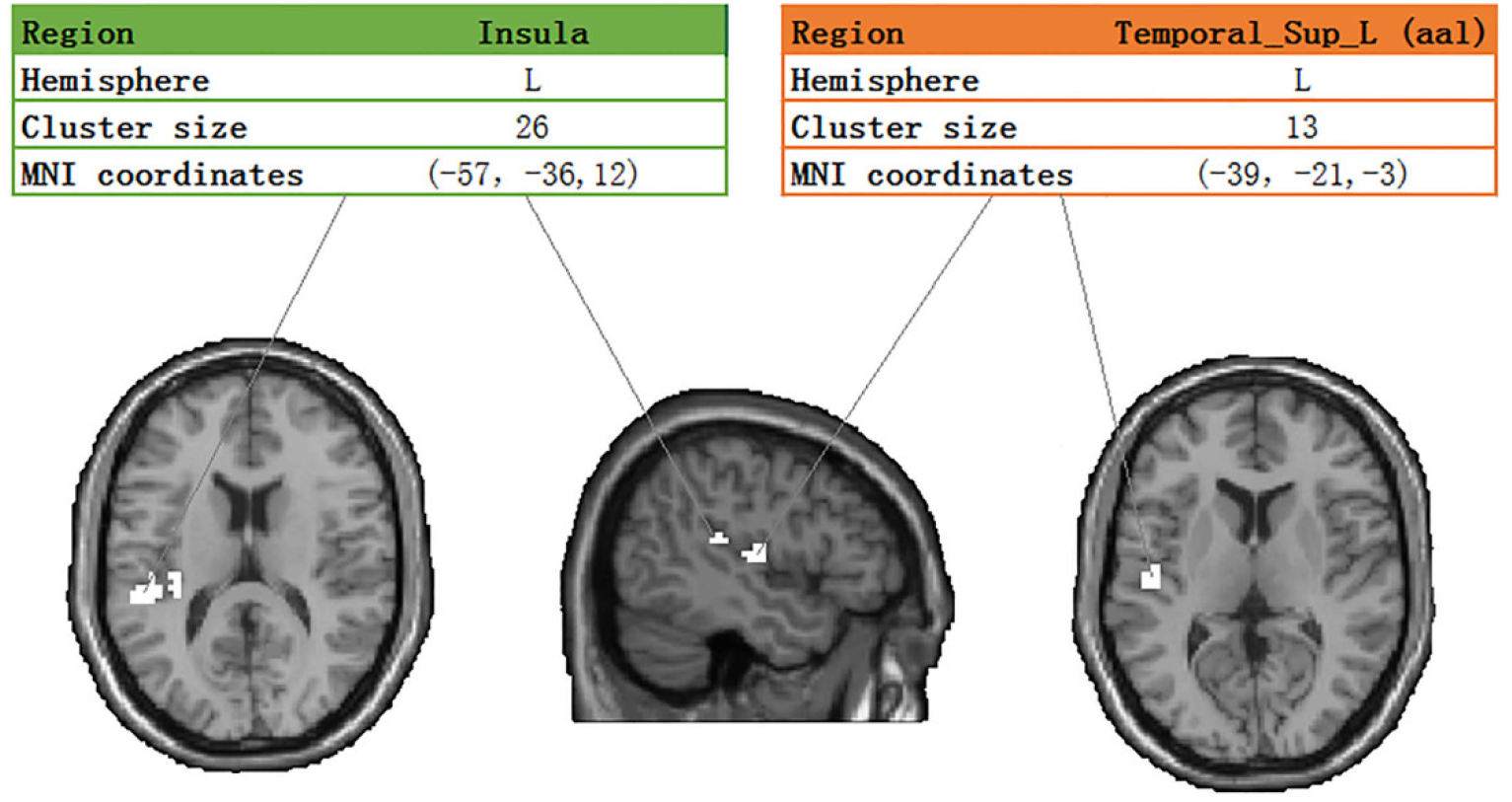
The conjunction analysis result of (Anger>Neutral)∩(Sad>Neutral)∩(Joy>Neutral). The cluster size indicated number of voxels; L, left; R, right.

## Discussion

In this study, the RSA based on ROIs, whole-brain searchlight analysis and emotion conjunction analysis were conducted to explore the brain regions involved in speech emotion perception in audio-visual modality. The results showed that the bilateral AMG was involved in the perception of positive and negative emotional stimuli, especially sensitive to negative emotions, but could not perceive neutral emotion. The left posterior insula and the left anterior STG were related to the perception of multimodal speech emotion in all valences, and bilateral IPL didn’t participate in the perception of neutral emotion.

Recently, there has been a study describes the variational implementation of covariance component analysis, which has the functionality of pattern component modelling (PCM) and RSA (Friston et al. 2019). It considers RSA and PCM as Bayesian model comparation procedures that assess the evidence for stimulus or condition-specific patterns of responses distributed over voxels or channels. In our study, we creatively proposed a weighted RSA method to evaluate the contribution of each candidate model to the best fitted model, which is also an enrichment and expansion of RSA.

### The perception of speech emotion in audio-visual modality

The RSA based on ROIs suggested that the left IPL, bilateral insula and bilateral STG were significantly associated with all single emotion models except the sad model. The searchlight analysis pointed out the significant correlation between the right IPL and neutral model and it was included to the brain regions that might be involved in the perception of emotion. Only the left posterior insula and left anterior STG were observed in the further conjunction analysis of emotion, indicating their roles in the perception of different speech emotion valences in audio-visual modality.

For a long time, the insula has been thought to process the physical sensations of appetite and disgust and arouse the associated emotions, leading to conscious perception of emotion states (Schachter and Singer 1962; Russell 2003). Evidence from electrophysiological studies and hemispheric inactivation processes suggested that emotional processing within the insula was strongly lateralized based on autonomous input from this region (Oppenheimer et al. 1992). For example, the bilateral insular cortex was activated (experience) when people smell unpleasant smells, but only the left side was activated (perceive) when they see other people doing the same thing, demonstrating the role of the left insula in perceiving emotions. A recent study showed that sensory testing of facial expressions was associated with the functional connectivity from the fusiform gyrus to the frontal lobe and left insular cortex, suggesting that the left insula might be involved in the perception of facial emotions (Bae et al. 2019). Other studies have found that the activation of emotion perception in the middle and posterior insula was biased towards the left side. These results were strongly consistent with our findings, which revealed the role of the left posterior insula in the perception of speech emotion and provided further evidence for the left lateralized of insula in emotion perception.

The STG was often revealed to be involved in the unimodal emotion perception (Dara 2012; Ethofer et al. 2012; Zhang et al. 2019). Previous studies have shown that the bilateral STG were significantly activated when hearing emotional sounds compared to neutral sounds (Ethofer et al. 2012), and that the right STG was associated with extracting tonal cues for emotional inferences, while the bilateral STG was involved in processing sonic cues of emotional prosody (Dara 2012). Studies also showed that the right STG played an important role in emotional facial perception of different valences (Zhang et al. 2019). Furthermore, the STG was also revealed to participate in processing of emotion perception in audio-visual modality. When the music and the face expressed the same emotion, the bilateral STG showed a stronger activation (Jeong et al. 2011). The bilateral anterior STG was observed in audio-visual emotion perception when using emotional prosody and dynamic facial expression as stimuli (Robins et al. 2009). A recent study used the meta-analysis to quantitatively summarize the results of 18 neuroimaging studies and identified a core processing network of emotion, including the right posterior STG, the left anterior STG, the right AMG and the thalamus, supporting the involvement of the STG in emotion processing of audio-visual modality (Gao et al. 2019). Although more and more studies have been conducted to explore the perception mechanism of emotion, the exact location related to emotion perception was not consistent in the results and most of the research pointed to the temporal lobe, usually biased to the right hemisphere (Robins et al. 2009). In this study, the short speech sentences with emotional information were used as the stimuli. However, previous studies on emotion perception mainly used music or emotional rhythm as auditory stimuli, and their conclusions might not apply to our findings. In order to eliminate the interference of semantic information in the study of emotion perception, the subjects were required to make emotion recognitions ignoring semantic information in sentences before the experiment, but this does not seem to work. Numerous studies have found that the left STG was involved in the processing of lexical and syntactic information, not the right side (Friederici and Kotz 2003; Pisoni et al. 2012; Feng et al. 2020). Our study indicated that the short speech sentences containing emotional information might subconsciously lead to the processing of semantic information in the brain, thereby excluding the right STG from the involvement in speech emotion perception, and provided evidence for the role of the left anterior STG in the perception of speech emotion.

### Regions that couldn’t perceive emotions of all valences

Compared with other emotion models, the bilateral AMG showed significant correlations with the negative emotion model in the RSA analysis. The weighted RSA suggested that the negative emotion model made more significant contribution to the best fitted model of bilateral AMG compared to other regions. These findings suggested that the AMG played a vital role in the perception of negative emotions. Previous studies have shown that the AMG was activated more significantly by negative emotional stimuli than neutral or positive ones (Irwin et al. 1996; Schneider et al. 1996; Lane et al. 1997; Zald and Pardo 1997), and the negative emotion processing of amygdala have been observed in both visual and auditory modalities (Schaefer et al. 2002; Koelsch et al. 2006), which were consistent with our findings. The amygdala might have a more general function in perceiving different emotional facial expressions (Fitzgerald et al. 2006), playing a central role in processing both negative and positive emotions (MORIOKA et al. 2010). In our study, the weighted RSA suggested that the anger and joy model made significant contributions to the best fitted model in bilateral amygdala, and statistical analysis revealed that the neural representation RDMs of bilateral amygdala were significantly correlated with anger and joy emotion models, which provided evidence for the amygdala in dealing with emotions of different valences. Based on the previous studies and our findings, we speculate that the amygdala may be involved in the representation of emotions in different valences, and is more sensitive to the perception of negative emotions. No significant correlations between the bilateral amygdala and neutral emotion model in the RSA based on ROIs and searchlight analysis were observed, and the amygdala was not detected in the conjunction analysis detect, indicating that the amygdala may not participate in the perception of neutral emotions.

The bilateral IPL was not observed in the further conjunction analysis, suggesting that the IPL might not be involved in the perception of speech emotion in audio-visual modality, which was consistent with our previous study that the IPL couldn’t decode emotion across modalities (Cao et al. 2018). Another study that explored the representations of emotions cross modalities suggested that the IPL could not classify the emotion of positive and negative valences, nor could it distinguish the emotional stimuli and neutral stimuli, which were similar to our results (Kim et al. 2017). We speculate the reason that the IPL was absent from the results of emotion perception might be that the IPL was unable to perceive the neutral emotion. Moreover, the neural emotion model made a quite small contribution in the weighted RSA. The statistical analysis showed that the correlation between the neutral emotion model and the neutral representation of right IPL did not reach the significant level, and the correlation value and significance value of the left IPL were both very low, which provided evidence for the incapability of bilateral IPL in perceiving neutral emotion. However, in our study, the searchlight analysis found a significant correlation between the bilateral IPL and the neutral emotion model, possibly due to the strong ability of RSA. When conducting RSA, the type I error might incorrectly infer the voxels that were not correlated to the model, thus obtaining the brain regions that shouldn’t exist (Thirion et al. 2015). Taking these into consideration, it’s considered that the bilateral IPL might not be associated with speech emotion perception in audio-visual modality.

### Emotion models

None of the ROIs showed a significant correlation with the sad emotion model, possibly because the perception of sad emotion was strongly influenced by individual differences. The individual differences have been suggested to be responsible for the inconsistencies in the findings of previous studies on sadness (Eugène et al. 2003). In our study, as the arousal of sadness was different among the subjects, all the ROIs failed to obtain a significant correlation with the sad emotion model at the group level. Other studies revealed that the spontaneous response of AMG was relatively low to sad faces (Kugel et al. 2008), and the people’s recognition ability to sad emotion decreased with age (Demenescu et al. 2015). These findings suggested that the brain’s ability of perception and recognition to sadness might be weak, and the perception difference between participants was relatively large, all of which might cause the insignificant correlation between the representation pattern of ROIs and the sad emotion model. In addition, our results indicated that the anger and joy emotion model were significantly correlated with neural representations of all ROIs and contributed prominently to the best fitted models. Some studies have found that excited and angry facial expressions could cause enhanced activations compared with neutral expressions, revealing an obvious “emotion effect”, which was consistent with our findings (Beltrán and Calvo 2015).

## Limitation

Emotion perception includes cognitive process and emotional process. The ROIs in this study are mainly concentrated in the ventral region of the brain, which is related to emotional process. However, what is the role of the dorsal regions located in hippocampus and anterior cingulate gyrus in speech emotion processing and whether they are involved in the integration of multimodal affective speech information are still unknown. Further research should be conducted in this regard. Moreover, the results have shown that the left posterior insula and the left anterior STG are involved in emotion perception, but whether they are associated with the decoding of emotion still needs verification. Further studies can be conducted to classify different affective stimuli in these regions to reveal their role in the recognition of emotion.

## Conclusion

In this study, we used dynamic facial expressions and affective speech to explore the brain regions that is involved in the perception of audio-visual emotional speech in different valences. Traditional RSA, a weighted RSA method, the whole brain searchlight analysis and the conjunction analysis of emotion were used for the analysis of experimental data. The results showed that the bilateral AMG was able to process positive and negative emotions, especially sensitive to negative emotions compared with other ROIs, but couldn’t process neutral emotion. The left posterior insula and the left anterior STG were related to the perception of multimodal speech emotion in all valences, and bilateral IPL didn’t participate in the perception of neutral emotion. In addition, the weighted RSA method is an enrichment and expansion of traditional RSA, which provides a new way to deal with neuroscience problems by using multiple models.

## Acknowledgement

This work was supported by the National Natural Science Foundation of China under Grant 61703302.

